# Rapid synaptic plasticity contributes to a learned conjunctive code of position and choice-related information in the hippocampus

**DOI:** 10.1101/2021.06.30.450574

**Authors:** Xinyu Zhao, Ching-Lung Hsu, Nelson Spruston

## Abstract

To successfully perform goal-directed navigation, animals must know where they are and what they are doing—e.g., looking for water, bringing food back to the nest, or escaping from a predator. Hippocampal neurons code for these critical variables conjunctively, but little is known about how this where/what code is formed or flexibly routed to other brain regions. To address these questions, we performed intracellular whole-cell recordings in mouse CA1 during a cued, two-choice virtual navigation task. We demonstrate that plateau potentials in CA1 pyramidal neurons rapidly strengthen synaptic inputs carrying conjunctive information about position and choice. Plasticity-induced response fields were modulated by cues only in animals previously trained to collect rewards based on these cues. Thus, we reveal that gradual learning is required for the formation of a conjunctive population code, upstream of CA1, while plateau-potential-induced synaptic plasticity in CA1 enables flexible routing of the code to downstream brain regions.

## Introduction

Animals rely on information from memories to perform many survival-related behaviors. In order to accurately encode experiences as unambiguous memories, information regarding the external environment and the animal’s internal behavioral states need to be bound together. A conjunctive code of positions and behaviorally relevant contexts can disambiguate representations of experiences that happened in a same spatial environment while the animal was engaged in different behaviors, thus providing critical information for memory-guided behaviors in the future. Here, we seek to understand how multiple dimensions are incorporated into conjunctive neural representations.

The hippocampus, a critical brain area for memory-guided behaviors (Squire, 2009; Morris, 1981), encodes both positional and behavioral information during navigation. Individual pyramidal cells in the hippocampus fire action potentials in specific regions of an environment (i.e., place fields) (O’Keefe and Dostrovsky, 1971); in addition, their firing is influenced by non-spatial factors, often referred to as behavioral context (Gulli *et al.*, 2020; Kentros *et al.*, 2004). This contextual coding means that hippocampal neurons are well suited for encoding memories of events that contain information about both the environment and the animal’s behavior. For example, a fraction of place cells in the CA1 subregion of the hippocampus has been shown to exhibit firing rate modulation based on the animal’s past and/or future trajectories in spatial alternation tasks (Wood *et al.*, 2000; Frank, Brown and Wilson, 2000; Ito *et al.*, 2015), which require animals to make alternating left and right turns in a figure 8-like maze, or to sequentially explore different arms of a W-shaped maze. In these tasks, some cells fire selectively in a common location prior to left or right turns. These so-called ‘splitter cells’ (Hasselmo, 2005) have also been observed in primates performing virtual navigation tasks (Gulli *et al.*, 2020). Following the original observations of splitter cells, numerous studies have found similar context-dependent firing in CA1 cells when the animal was involved in various behavioral tasks, including spatial alternation tasks with a delay period between each trial, implemented using a holding chamber or running wheel (Pastalkova *et al.*, 2008; Wang *et al.*, 2015; Ainge *et al.*, 2007b), and cued licking behavior in olfactory association tasks with a delay in head-fixed animals (MacDonald *et al.*, 2013). It has been suggested that context-dependent firing, as demonstrated in splitter cells, is crucial for unambiguously encoding specific spatial memory episodes and, consequently, guiding behaviors (Hasselmo and Eichenbaum, 2005; Ainge *et al.*, 2007b; Ainge *et al.*, 2007a; Ito *et al.*, 2015; Kinsky *et al.*, 2020). These complex tuning properties of neurons are believed to form during learning. Despite their conceptual importance, the plasticity mechanisms underlying the formation of splitter cell activity in CA1 remains unclear.

Diverse forms of synaptic plasticity have been reported in CA1 (Malinow and Miller, 1986; Remy and Spruston, 2007; Dudman, Tsay and Siegelbaum, 2007; Bi and Poo, 1998; Bittner *et al.*, 2017; Kim *et al.*, 2015), but it remains unknown which of these is responsible for forming conjunctive hippocampal representations. Recently, it was shown that plateau potentials, a form of large-amplitude, long-duration calcium spikes, can rapidly induce a robust synaptic potentiation—behavioral time-scale synaptic plasticity (BTSP)—in CA1 pyramidal cells (Bittner *et al.*, 2015; Bittner *et al.*, 2017). CA3-to-CA1 synapses that were active within 1-2 seconds relative to a given postsynaptic plateau potential can be effectively potentiated. Several lines of evidence suggest that BTSP plays an important role during rapid establishment of feature selectivity in CA1 pyramidal cells. First, BTSP can potentiate synaptic inputs even with a single plateau potential, suggesting it as a powerful mechanism for rapid learning (Bittner *et al.*, 2015). Second, BTSP naturally converts silent cells into active, spatially tuned place cells (Bittner *et al.*, 2015). Third, BTSP can create place cells that distinguish between identical visual cues presented within different environments (Zhao *et al.*, 2020). These observations collectively implicate BTSP as a physiological mechanism for flexibly supporting a conjunctive code in the hippocampus. However, it remains unknown whether splitter cells could be generated through rapid plasticity induced by plateau potentials in memory-based spatial tasks.

To address this question, we performed whole-cell patch-clamp recordings in head-fixed mice running in a virtual-reality (VR) system, which facilitated precise control of sensory stimulation and intracellular recording and manipulation of single cells. We previously demonstrated that population activities of CA3 cells can be inferred from the membrane potential dynamics in BTSP-induced CA1 place fields (Zhao *et al.*, 2020). Therefore, intracellular recordings and manipulation of CA1 pyramidal cells, which allows triggering of Ca^2+^ plateau potentials, provides a unique opportunity for testing the role of cellular plasticity mechanisms as well as assessing the nature of the presynaptic inputs for the conjunctive code at the same time. In the current study, mice were trained to run down a linear track and turn into one of the two choice arms from a Y-junction to collect rewards, based on distinguishable cues visible only at the very beginning of the track (*initial cues*) but not at the choice point. Our results showed that plateau potentials induced trial-type-specific, spatially tuned firing patterns in almost all recorded CA1 cells. The differential firing of induced splitter cells was primarily attributable to a trial-type-specific synaptic depolarization that was only observed in mice trained to perform the cued two-choice turning task. Together, our results reveal that the rapid synaptic plasticity induced by plateau potentials can support the manifestation of a conjunctive code of the animal’s current position and task/choice-related information in CA1, thus enabling the flexible relay of task-specific variables to extra-hippocampal brain regions.

## Results

### The dorsal hippocampus is required for mice performing cued Y-maze tasks in virtual reality

Adult mice were head-fixed and trained to run on a spherical treadmill. Visual scenes were rendered on three monitors, with imagery covering a large portion of the animal’s visual field (217° in azimuth and 80° in elevation) and the optic flow coupled with the animal’s locomotion (Cohen, Bolstad and Lee, 2017). Mice were trained to run unidirectionally on a linear corridor and turn into one of the two choice arms at a Y-junction to collect rewards. The turn direction was cued by distinguishable visual cues at the first 45 cm of the corridor (“initial cues”, Fig. 1A). Within this “cue zone”, high-contrast vertical gratings and circles were presented on the two separate sides of the walls, with the vertical gratings always associated with the rewarded side. Following the cue zone, up to the choice point (Y-junction) and choice arms, cross patterns were painted on both sides of the walls in a 95-cm “delay zone”; it provided optic flow during the animal’s running, without offering any information about the reward location. At the end of each trial, the animal was teleported back to the beginning of the Y-maze for the next trial. Cued directions were randomized from trial to trial (see Methods for details). All mice used in our analyses went through a progressive training curriculum (Supplementary Fig. 1), and eventually performed the final task with a >80% success rate (on average, ~90%) before being used in behavioral or physiological experiments.

**Figure 1.**
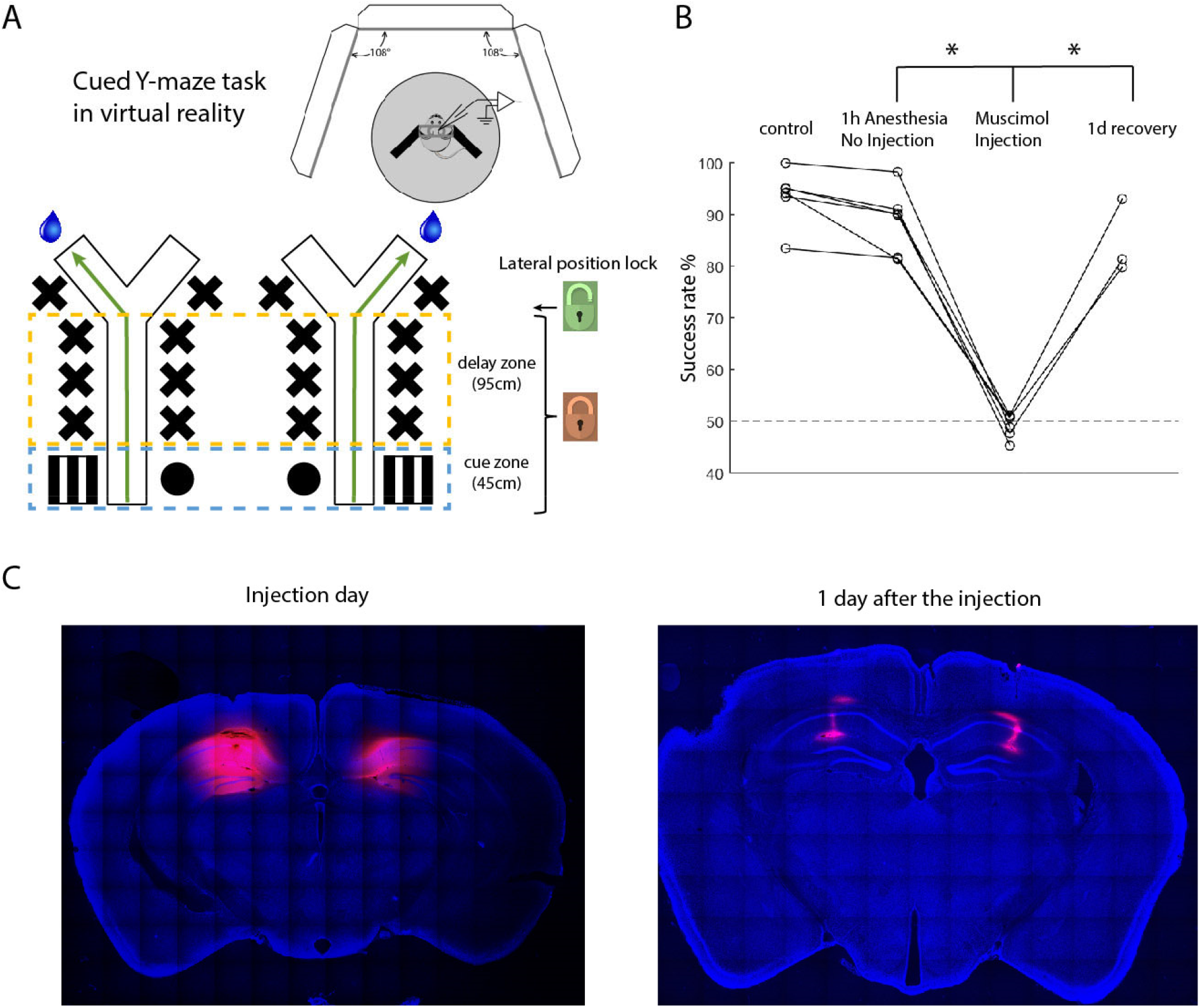
The dorsal hippocampus was required for mice to perform the cued Y-maze task in virtual reality. **A.** Experimental setup. Head-fixed mice running on a spherical treadmill were trained to collect rewards in a two-alternative, cued-Y maze task in virtual reality. The animal’s lateral position was locked on the midline and the view angle remained straight (the brown closed lock symbol) until it reached the Y-junction (the choice point; the green open lock symbol). **B.** Bilateral silencing of the dorsal hippocampus diminished the performance to the chance level in trained animals. A total of 6 mice were tested in the task after the injection of 3.2mM BODIPY TMR-X muscimol conjugate. 3 of them were sacrificed immediately after the behavioral test for histological examinations, whereas the others were tested again one day after the injection (p=5.576e-6 for the muscimol injection group vs. 1h-anesthesia-only group, n=6 mice; p=0.0115 for the muscimol injection group vs. 1d-recovery group, n=3 mice; two-tailed paired Student’s t-test). *: significant difference, p<0.05, paired student t-test. **C.** Coronal sections of the brain confirmed that the drug injection was confined within the dorsal hippocampus with good coverage (red: TMR-X muscimol conjugate; blue: DAPI). A small patch of the fluorescent signal was only visible around injection site one day after the injection.

In the delay zone of the Y-maze task for freely moving mice/rats, differential sensory inputs would be introduced due to potential behavioral or gait biases (e.g., the body running closer to either side or the head pointing toward either direction) in left- vs. right-rewarded trials even though identical visual landmarks were present. As a result, it may confound the interpretation of differential neural activities observed during the delay zone as truly behavioral context dependent since an alternative explanation for them is an immediate response to distinct visual inputs. This issue can be addressed with virtual reality: we locked the animal’s virtual position along the midline, with no degree of freedom for the virtual head direction before the animal reached the Y-junction. At the choice point, the animal could bias the treadmill rotation to choose which arm to turn into. Therefore, in our behavioral design, the gross visual inputs during the delay remained identical regardless of the animal’s behavioral biases.

To test whether the dorsal hippocampus is involved in performing the task, we silenced it bilaterally by injecting fluorescently conjugated muscimol (Allen *et al.*, 2008) in well-trained mice. *Post-hoc* histology confirmed that the drug was confined within the targeted area (Fig. 1C). 1-hour general anesthesia, as a sham operation, did not affect the performance. Silencing the dorsal hippocampus dramatically reduced the success rate in the Y-maze, from 88.7 ± 2.6% to 49.0 ± 0.9% correct (mean ± S.E.M. and hereafter), which is around the chance level (Fig. 1B). Performance recovered one day after the injection (84.7 ± 4.2%, Fig 1B), consistent with the observation that the fluorescence disappeared from most of the hippocampus (except for a narrow track close to the injection sites, Fig. 1C). The strong and fully reversible behavioral effects of pharmacological inactivation demonstrated that the intact function of the dorsal hippocampus is required for the performance in the cued Y-maze task, after the mice have learned it.

### Ca^2+^ Plateau potentials potentiated choice-dependent synaptic input in the delay zone

We next sought to exam functional properties of CA1 place fields induced by the rapid synaptic plasticity in this memory-guided spatial navigation task. Place cells have been shown to be induced by postsynaptic Ca^2+^ plateau potentials, via synaptic plasticity, occurring either spontaneously or experimentally (Bittner *et al.*, 2015). To test the effects of triggering this form of plasticity in our paradigm, intracellular whole-cell recordings were made from CA1 pyramidal cells by standard blind patch methods (Zhao *et al.*, 2020) (see Methods). All cells included in this dataset did not show any firing fields prior to the induction. Plateau potentials were triggered by brief current injections (700-900 pA, 300 ms) when the animal arrived at an experimentally defined position in the delay zone before the junction (110 cm from the beginning of the track, Fig. 2A). Importantly, these current injections were performed only on selected trials (1-7 trials in total) where the animal was cued to a specific direction (referred to as “plasticity direction” or “P-direction” hereafter), which was chosen randomly for each experiment; no current was delivered on trials cued for turns in the opposite direction (referred to as “opposite direction” or “O-direction” hereafter). In our 14 stable recordings, left-cued and right-cued trials were selected for the P-direction in 6 and 8 experiments, respectively. We found that this induction protocol resulted in robust action-potential firing peaked around the position where the current injection was applied (Fig. 2A, B). These induced response fields, presumably driven by potentiated synaptic inputs (Bittner *et al.*, 2017), persisted throughout the entire recording session without any further experimental current injection. Remarkably, most of the cells tested showed spatially modulated fields with the property consistent with splitter cells: 86% of the cells (n=12/14 cells from 6 mice) exhibited the normalized difference of peak firing rates between the two turn directions larger than 70% (see Methods; Fig. 2C). In fact, 10 cells showed a peak firing rate difference close to 100%, meaning they fired almost exclusively in P-direction trials.

**Figure 2.**
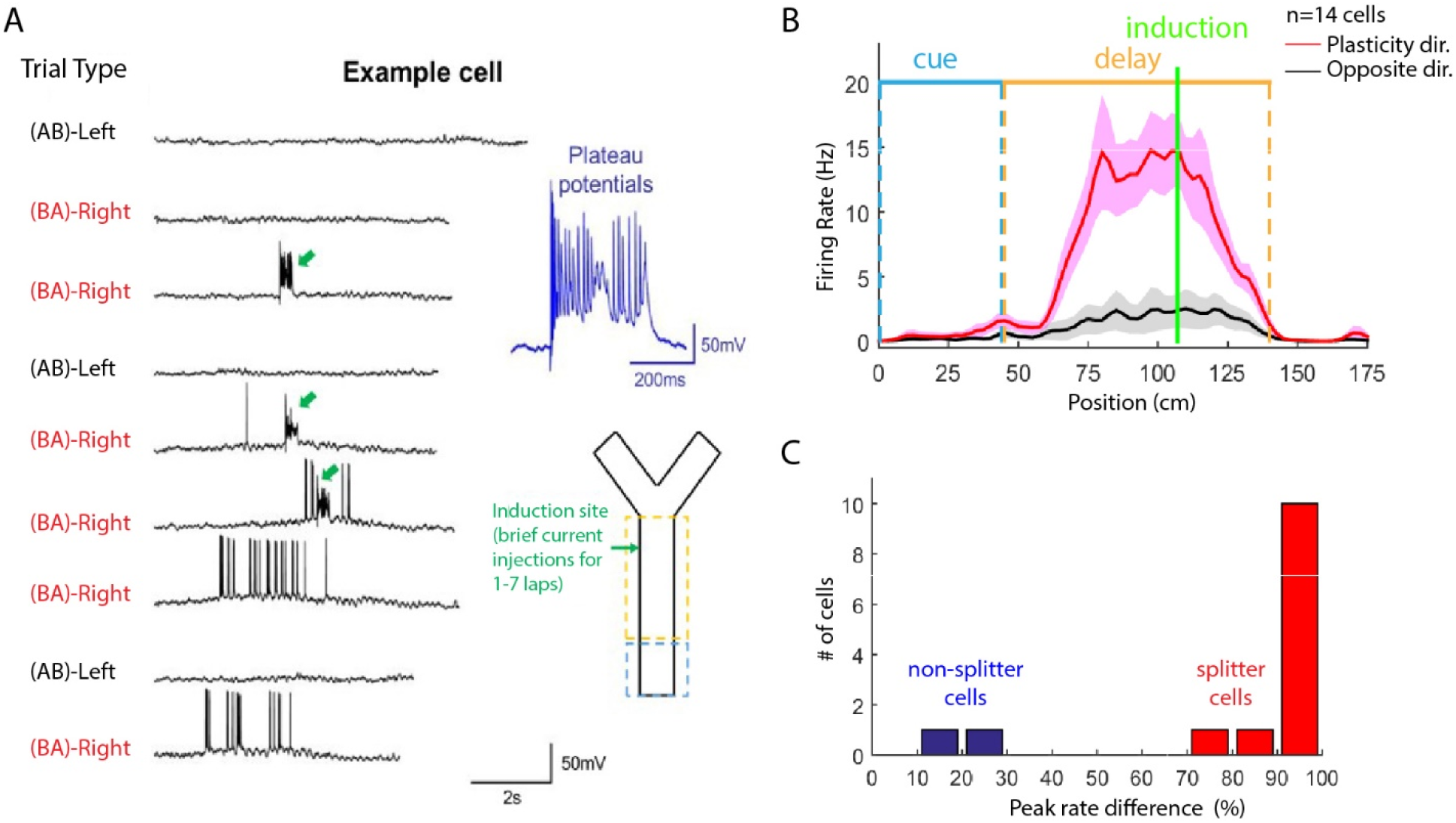
Rapid synaptic plasticity induced trial type-specific activities in most of the tested CA1 pyramidal cells in mice performing the cued Y-maze task. **A.** Place-field induction by the rapid plasticity in an example cell. Traces starting from the top are whole-cell recoding from consecutive trials, with the trial type indicated on the left. Green arrows indicate plateau potentials triggered by the large, brief current injections (inset, *top-right*: the plateau potential from the first current injection). Upon the third induction in the (BA)-Right trial, a firing field had emerged and exhibited only in (BA)-Right trials. Inset, *Bottom-right*: schematic of the Y-maze, with the location where the brief somatic current injection indicated (green arrow). **B.** Averaged firing rates at different positions after the plasticity induction (red: plasticity direction, black: opposite direction, n=14 cells from 7 mice). Prior to the induction, all the neurons tested were silent cells. Line and shades represent mean and S.E.M. Blue and yellow bars mark the range of cue and delay zones, respectively. **C.** Distribution of cell number by the normalized peak firing rate difference after the plasticity induction. Peak rate was calculated from the averaged curve of the spatially tuned firing rates across trials.

Running speed, which has been reported to modulate the activity of CA1 cells (Fuhrmann *et al.*, 2015), and the associated visual inputs were not the determinant for the choice dependency of the splitter cells. Neither the animal’s locomotion speed nor the optic flow speed in VR was significantly different between different trial types (Supplementary Fig. 2).

It should be noted that some of the trials in which the current injections were applied could be error trials as even well-trained animals made a small number of mistakes (e.g., it was cued to the left but turned to the right). For both non-splitter cells, the animals made some errors during the induction trials (cell #1 and #2, Supplementary Table 1). We do not rule out the possibility that these error trials contributed to the activity of these cells in trial-types where plateau potentials were not induced. However, error trials during induction did not prevent the formation of a splitter-cell response (cell #5, #6, #10, and #13, Supplementary Table 1). Two possibilities may explain this observation: (1) Although in some cases, triggering plateau potentials in only 1 or 2 trials was sufficient to induce a significant response field, most cells required more. Since there were more correct trials (and only 1 or 2 error trials) during induction for all recorded cells, it is possible that any effect of error-trial induction could not be a factor for dominating the property of the induced fields. (2) The hippocampus sometimes still encodes the cue configuration correctly even when animals made wrong choices (i.e., hippocampal representations were dominated by cues but not motor patterns), as will be shown in a later section.

Given that plateau potentials efficiently generated context-dependent place-cell firing likely through rapid potentiation of presynaptic inputs, the integrated postsynaptic response of splitter cells may provide interesting clues regarding the properties of the inputs after the task has been learned. Four possible input patterns may lead to the context-dependent firing (Fig. 3A, *from top to bottom*): (1) Initial cues may trigger a prolonged place field in one of CA1’s input circuit, which is later integrated with the positional input from another pathway. (2) Slow, position-invariant, contextual inputs may integrate with a positional input, showing a *DC*-like membrane potential offset in different trial types. (3) The contextual signal may exert multiplicative effects on a context-independent positional signal and thus act as gain control. (4) Separate representations of the two trial types may be already formed in a major presynaptic input area to CA1, so that potentiation of the synapses active in one trial type will have little effect on the synapses active in the other trial type.

**Figure 3.**
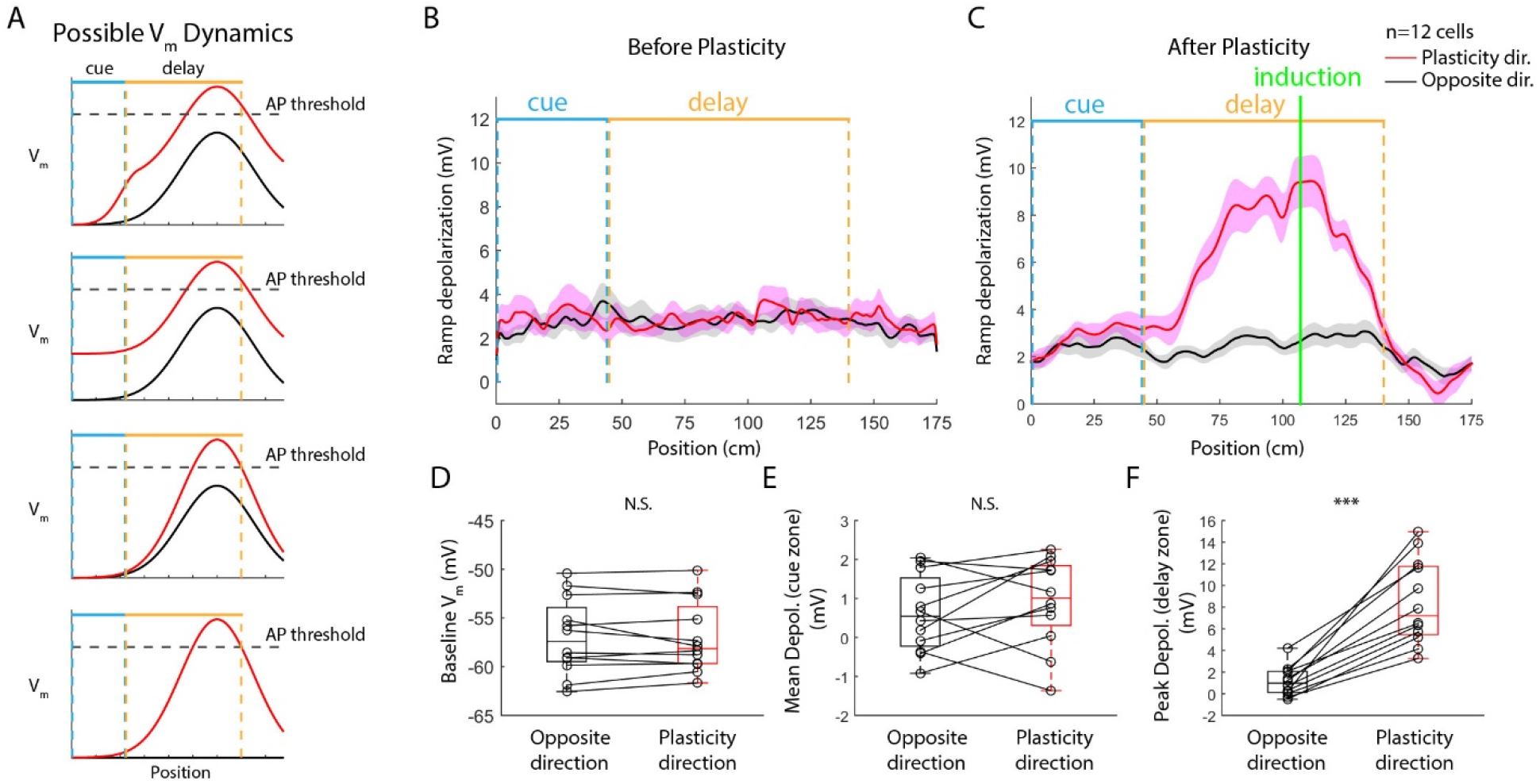
Plasticity induced differential depolarization in the delay zone without altering the baseline membrane potential or the depolarization in the cue zone between different trial types. **A.** Possible membrane potential dynamics underlying the trial-type-specific firing pattern. **B and C.** Averaged Vm ramp depolarization before (B) and after (C) the plasticity in the induced splitter cells (red: plasticity directions, black: opposite direction, n=12 cells from 6 mice). Curves and shades represent mean and S.E.M. Blue and yellow lines mark the cue and delay zones, respectively. **D, E and F.** Statistical comparisons of the baseline V_m_ (D), cue-zone depolarization (E) and delay-zone depolarization (F) between plasticity and opposite directions after the plasticity induction (p=0.7943 for the V_m_ baseline; p=0.2923 for the mean depolarization in the cue zone; p=1.242e-5 for the peak depolarization in the delay zone; n=12 cells from 6 animals; two-tailed paired student t-test). N.S.: non-significant, p>0.05. ***: significant difference, p<0.001. All metrics were calculated after averaging across trials in each recorded cell.

In order to distinguish these possibilities, we investigated the subthreshold membrane potential (V_m_) underlying the context-dependent firing of these 12 induced splitter cells. We found that (Fig. 3 B, C) V_m_ baseline and depolarization in the cue zone did not significantly differ between the two turn directions (V_m_ baseline: −57.0 ± 1.1 mV for plasticity direction vs. −56.9 ± 1.1 mV for opposite direction, Fig. 3D; cue zone depolarization: 0.9 ± 0.3 mV for plasticity direction vs. 0.6 ± 0.3 mV for opposite direction, Fig. 3E). However, a significantly larger difference in depolarization was observed in the delay zone for the two turn directions (8.4 ± 1.1 mV for plasticity direction vs. 1.1 ± 0.4 mV for the opposite direction, Fig. 3F). Besides, the turn direction-specific ramp depolarization was accompanied by an increased theta oscillation amplitude (3.8 ± 0.5 mV for plasticity direction vs. 1.3 ± 0.1 mV for opposite direction, Supplementary Fig. 3), which is similar to what has been found for the synaptic ramps in previous studies (Bittner *et al.*, 2015; Zhao *et al.*, 2020; Harvey *et al.*, 2009).

Remarkably, we found that each silent cell could be turned into a splitter cell with trial-type specificity determined by arbitrary selection of a plasticity direction for induction. The induced response is all-or-none like (Fig. 3C), suggesting that the presynaptic position-dependent inputs activated in the two trial types might be separate, non-overlapping populations. Consistent with this idea, in one recording, we first induced a field in left-cued trials, followed by an induction in right-cued trials (Supplementary Fig. 4). We found that the second induction resulted in a right-preferring field with the response size comparable to the left-preferring field, without affecting much the previously induced field. The lack of occlusion effects further supports the notion of separate groups of activated presynaptic cells in different trial types.

In addition to potentiating excitatory inputs from upstream areas, plasticity can also alter the inhibitory input from local interneurons. A recent study showed that stimulation of CA1 pyramidal cells can remap place fields through plasticity between pyramidal cells and interneurons (McKenzie *et al.*, 2021). To test whether the induced trial-type specificity may originate from local interneurons, we identified 6 putative fast-spiking interneurons in our recordings by their narrow spikes and relatively high firing rates, as shown in an example recording in Supplementary Fig. 5A. Consistent with previous studies (Góis and Tort, 2018), the cell is highly modulated by the animal’s locomotion. Interestingly, the cell showed significant spatial tuning, with higher firing rates in the cue zone and the choice (reward) arms, compared to the delay zone. Such a spatial profile was also observed on the population level in terms of averaged membrane potential depolarization (Supplementary Fig. 5B) and firing rate (Supplementary Fig. 5C), which appeared consistent among all 6 recorded cells (Supplementary Fig. 5D). Previous studies in freely moving rats during random foraging have shown poor spatial tuning of fast-spiking interneurons in CA1 (Wilson and McNaughton, 1993). The reason for the emergence of this spatial pattern in interneurons recorded in our Y-maze task remains to be investigated in future. One possibility is that the CA1 population over-represents the reward location, as shown in previous studies (Gauthier and Tank, 2018), and probably also the cue zone (Sato *et al.*, 2020), which is reward-predicting. Fast-spiking interneurons may thus track the overall firing rates of pyramidal cells to balance the network activity, as shown previously in both CA1 and neocortical areas (Grienberger *et al.*, 2017; Atallah *et al.*, 2012; Liu *et al.*, 2009). Nonetheless, putative fast-spiking interneurons were only moderately modulated by trial type (Supplementary Fig. 5E). Even if plateau potentials also modify the synapses between pyramidal cells and interneurons, the weak trial-type specificity in interneurons makes them unlikely to be the primary factor that led to the large trial-type-dependent depolarization induced by plateau potentials (Fig. 3). Furthermore, the interneuron-mediated plasticity often generated responses at locations unrelated to where the manipulation was done (McKenzie *et al.*, 2021), while in our recordings the plasticity induced place fields always peaked around the location where plateau potentials were triggered. These results are consistent with a parsimonious explanation that the rapid induction of trial-type specific depolarization by plateau potentials is primarily mediated by potentiating excitatory synapses.

In summary, our results demonstrate that plateau-potential-induced synaptic plasticity can rapidly and efficiently produce context-dependent CA1 place fields in mice performing a two-alternative cued Y-maze task. Furthermore, each silent CA1 neuron can be converted to a splitter cell with an arbitrary trial-type preference. Therefore, after the task has been learned, the presynaptic circuits might support the flexibility of rapidly switching the function of CA1 cells from silent to context-dependent place cells.

### Splitter cell activity correlates with either past visual cues or choice-related factors in error trials

There are multiple factors that may be crucial for the modulation of place-field responses by the trial type, including initial cues and the animal’s choice-related behavior. When the animal makes a correct choice based on the initial cues, these two factors are correlated. On the contrary, error trials, in which these two factors dissociate, provide an opportunity to assess their relative contribution to the synaptic response.

Individual mice developed unique running patterns to perform the task after training (Supplementary Fig. 6). Some mice quickly tilted their bodies toward the cued direction from the beginning, while the others held the same running posture for a certain distance before diverging to differential behavior. On average, when sorted by turn direction, their locomotion direction (indicated by the lateral running speed) exhibited small biases from the very beginning (even in the cue zone), which gradually increased throughout the delay zone until the choice point (Fig. 4A). On the other hand, the forward running speeds, which controlled the speed of VR optic flow, did not differ (Fig. 4B).

**Figure 4.**
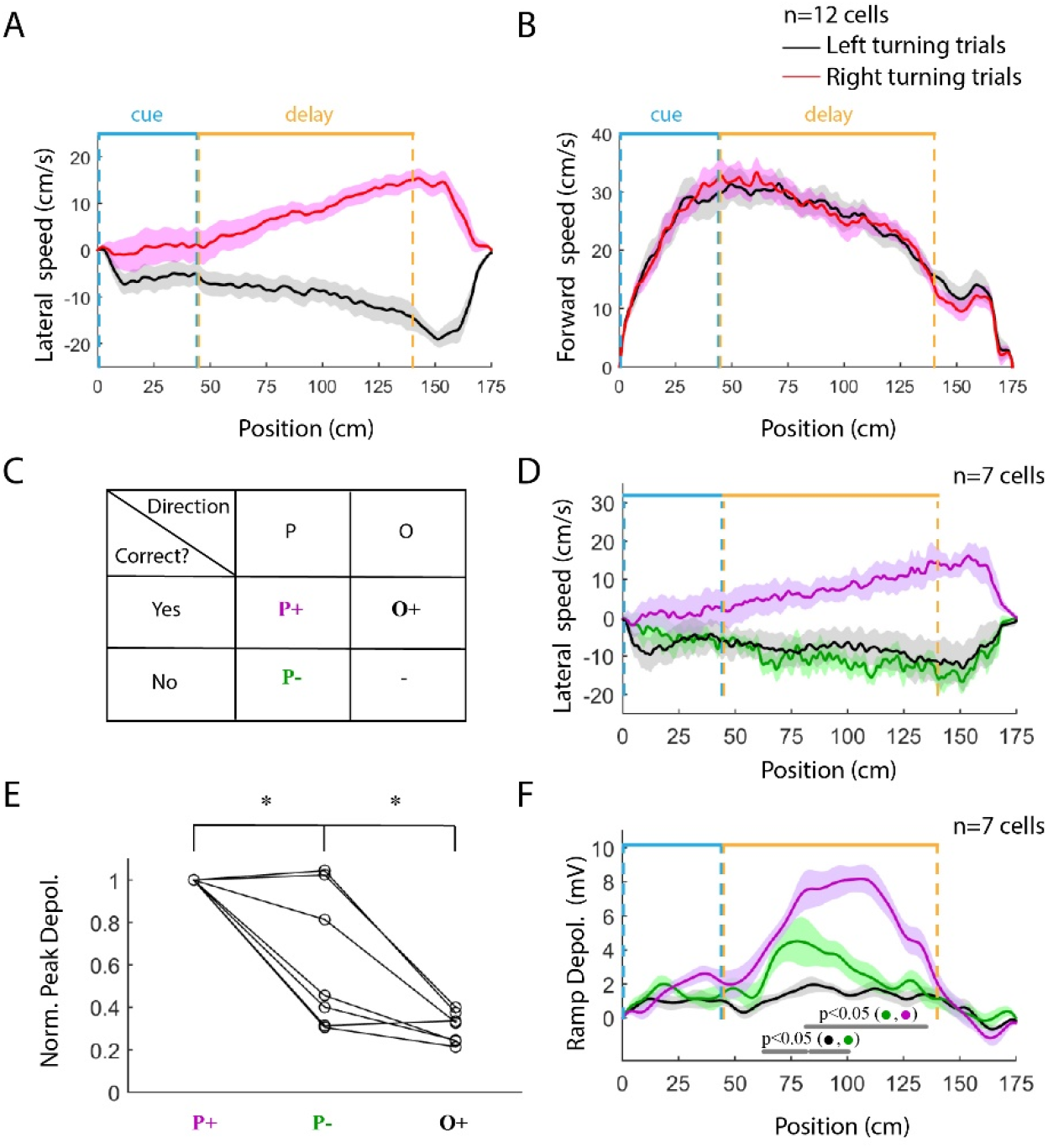
Both the past visual cues and the choice-related factors were reflected in the differential synaptic response in the delay zone. **A, B.** Averaged forward (A) and lateral (B) running speed. Red: left-turning trials; Black: right-turning trials. Curves and shades represent mean and S.E.M (n=12 recording sessions from 6 mice). **C.** Definitions of trial types. P and O indicate if the correct direction is the same as the P+: plasticity direction, correct trials; P-: plasticity direction, incorrect trials; O+: opposite direction, correct trials. O-trials (opposite direction, incorrect trials) were not included in the analysis because they were rare. **D.** Average lateral running speed in P+ (purple), P-(green), and O+ (black) trials. Only 7 cells from 5 mice were included in the analysis due to the small number of incorrect (P-) trials. **E.** Statistical comparison of the normalized peak depolarization in the delay zone between the three trial types (p=0.0229 for the P+ vs. P-trials; p=0.0221 for the P- vs. O+; two-tailed paired student t-test; n=7 cells from 5 animals). Ramp depolarization was calculated after averaging across trials and normalized to the P+ trial value. *: significant difference, p<0.05. **F.** Average ramp depolarization in P+ (purple), P-(green), and O+ (black) trials. Grey bars at the bottom of B represent positions where the green curve is significantly different from the purple and black curves, respectively (p<0.05 with two-tailed paired student t-test; n=7 cells from 5 mice). Curves and shades represent mean and S.E.M. Blue and yellow lines mark the cue and delay zones, respectively.

We analyzed the plasticity-induced response field in 7 cells from the mice that made relatively more errors (i.e., cued to one direction but turned to the opposite direction). Running behavior for correct and incorrect trials for the P-direction (“P+” and “P–” trials, respectively) was compared to correct trials for the O-direction (“O+” trials) (Fig. 4C; the incorrect trials for the O-direction were not included for the analysis due to the low trial number), and we found the running behavior was similar on P– and O+ trials (Fig. 4D), as expected. However, the intracellular recordings during these error (P-) trials exhibited diverse results, with the peak depolarization similar to that in P+ trials in some cells while similar to O+ trials in the other cells (Fig. 4E). On the population level, the ramp depolarization for P– trials was significantly different from both P+ trials and O+ trials (peak ramp depolarization: 9.0 ± 0.9 mV for P+ trials; 5.5 ± 1.3 mV for P– trials; 2.7 ± 0.4 mV for O+ trials; Fig. 4E and 4F), suggesting that both the past visual cues and the choice-related factors may each contribute to the differential inputs for the plasticity-induced fields.

Our analysis did not distinguish between multiple choice-related factors, including perception of visual cues, decision making, goal representation, efference copy of motor commands, and proprioception. Future work is needed to determine the relative contribution of these factors.

### Cue-reward association is required for initial visual cues to modulate the plasticity-induced place field in the delay zone

We next investigated whether the influence of the initial visual cues on the induced response field in the delay zone relies on their ability to elicit the differential behavior.

To address this question, we trained another set of mice without a fixed relationship between reward delivery arm and the initial cues. In this version of the task, distinct visual cues were still presented in the initial zone and varied randomly trial-by-trial, as described before, but no longer predicted the reward in any way. Thus, the mice had been exposed to training environments identical to the original task, but did not actively follow specific rules associated with initial cues anymore. For the 7 cells tested under this condition, plateau potentials were triggered in trials of only one cue configuration, but they generated nearly identical place fields between trials of both cue conditions (peak ramp depolarization in the delay zone: 8.8 ± 1.1 mV for plasticity cue config. vs. 7.6 ± 1.2 mV for opposite cue config., Fig 5). Notably, since the mice were trained in a scheme that did not require specific turning to enhance the reward probability, there was no motivation to modify their innate biases related to running gait. As a result, in 6 out of 7 recordings, the mice only ran in one direction, which partially explains why the induced fields showed no context dependence about the initial cues. However, we argue that the lack of differential behavior cannot fully account for the results for the following reasons: first, our previous analysis of error trials suggested that the initial visual cues might affect later responses, even when dissociated from the correct behavioral choice (Fig. 4); second, in one experiment in which the mouse turned to more than one direction, the response amplitudes were only moderately different between left- vs. right-turning trials (Supplementary Fig. 7), which goes against the possibility that the homogeneous motor actions are the only factor determining the induced response dissociated from the context of the initial cues. Finally, it is possible that the information related to the initial visual cues is not transferred to the dorsal hippocampus at all due to the lack of behavioral importance. It is not the case, however, given that in 2 recordings, we observed synaptic responses starting in the cue zone that clearly differentiated the two distinct cue configurations (Supplementary Fig. 8). Together, these results imply that the initial sensory cues influence induced hippocampal place fields in the following delay zone only when the meaning of the cues has been learned as a relevant context for the task. It is possible that the presynaptic circuits in the hippocampus and/or associated brain areas are modified as the animal learns how to use cues to predict rewards, which allows the rapid plasticity to serve as a robust cellular mechanism for flexible routing of the context-dependent information to downstream regions.

**Figure 5.**
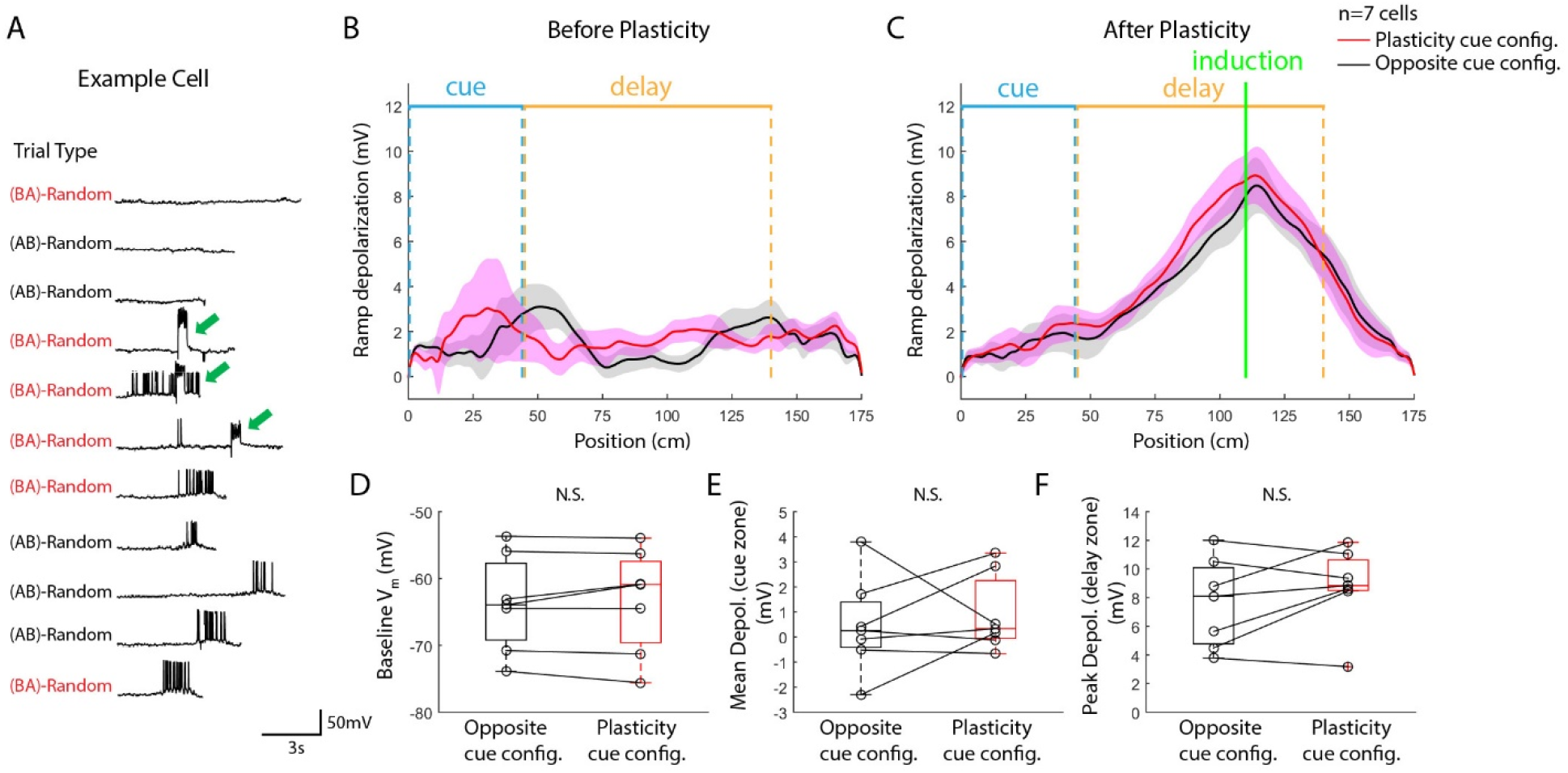
The rapid plasticity induced in trials of one cued condition resulted in nearly identical place fields in trials of both cued conditions when the initial cues were not associated with reward location during training. **A.** Place-field induction by the plasticity in an example cell from a mouse trained with random reward delivery. Green arrows indicate plateau potentials triggered by large, brief somatic current injection. In this cell, the plateau potentials in trials with the (BA) initial cue configuration induced place fields in both trial types. **B, C.** Averaged Vm ramp depolarization before (B) and after (C) the plasticity. (red: plasticity cue configuration, black: opposite cue configuration, n=7 cells from 5 mice). Curves and shades represent mean and S.E.M., respectively. Blue and yellow bars mark the cue and delay zones. **D, E, F.** Statistical comparisons of the baseline Vm (D), cue-zone depolarization (E) and delay-zone depolarization (F) between two cue configurations after the induction of plasticity (p=0.6093 for the Vm baseline; p=0.5776 for the mean depolarization in the cue zone; p=0.2095 for the peak depolarization in the delay zone; n=12 cells from 7 mice; two-tailed paired student t-test). Note: 1.5 X IQR (inter-quartile range) was chosen for the whisker plot, but all the data points were included for statistical comparisons; no points were removed as outliers. N.S.: not significant, p>0.05. All metrics were calculated from averages across trials in each recorded cell.

## Discussion

By obtaining intracellular recordings in mice performing a learned behavior in virtual reality, we investigated a cellular plasticity mechanism that supports the manifestation of a conjunctive code of the animal’s current position and choice-related information in the CA1 region of the hippocampus. Three key observations were: (i) Rapid synaptic plasticity triggered by plateau potentials can efficiently induce context-dependent place fields in CA1 cells, when the animal is performing well on a hippocampus-dependent, two-choice navigation task. (ii) The induced context-dependent firing is driven by a synaptic response (V_m_ ramp depolarization) largely restricted to the delay zone—the location of plasticity induction—where behaviorally relevant sensory cues are no longer visible to the animal. (iii) A fixed cue-reward association, learned by the animal during training, is required for the initial cues to modulate the induced place field in the delay zone.

Our findings demonstrate that the rapid plasticity induced by plateau potentials (i.e., BTSP), previously shown to naturally produce new place fields (Bittner *et al.*, 2015), can convert silent CA1 cells into splitter cells, which encode conjunctive information of the animal’s position and choice-related behavior. One prominent feature of BTSP-induced response fields is that for a given cell, the context-specific depolarization could be introduced to an arbitrary trial type, but the induced depolarization is almost absent in the opposite trial type. The most straightforward explanation is that most silent cells receive some weak and balanced inputs that already have strong spatial tuning *as well as* distinct tuning for the two trial types (Fig. 6). The set of synapses that gets potentiated is determined by the animal’s position and choice when plateau potentials are triggered in the postsynaptic cell. Separate representations of different trial types are likely formed during the training process with the fixed cue-reward association, as induced place fields in mice trained with random reward delivery did not show trial-type specificity. It should be noted that how individual presynaptic cells in CA1’s upstream areas change their firing properties during training remains an open question.

**Figure 6.**
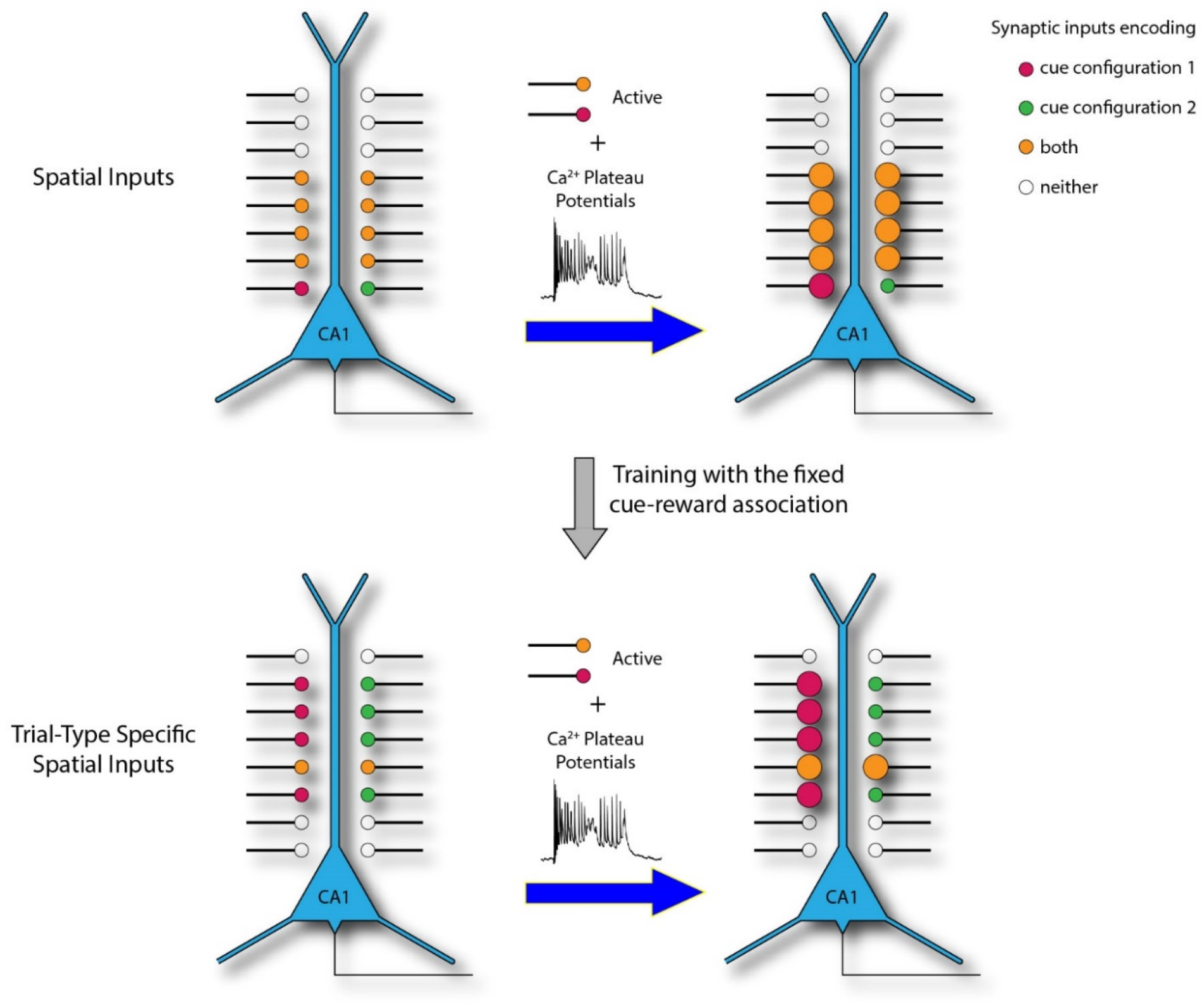
Working model. Behaviorally relevant sensory cues drive network activity in CA1’s presynaptic areas into different states. These states are presumably supported by largely segregated subpopulations of presynaptic neurons (represented by red and green axons/synapses), which have been formed during the training that associates the initial cues and the reward. Without training with the fixed cue-reward association, most inputs are active in trials with both cue configurations (orange). Plateau potentials can rapidly and efficiently potentiate the synapses activated during a given trial type and induce the formation of splitter cells.

One brain area that can provide such a conjunctively tuned input is CA3, which is a major excitatory driver of CA1 (Davoudi and Foster, 2019). Previous physiological (Lee *et al.*, 2004; Wills *et al.*, 2005) and theoretical (Rolls, 2007; Marr, 1971; Solstad, Yousif and Sejnowski, 2014) studies have suggested that CA3 circuits exhibit attractor dynamics arising from recurrent connections between pyramidal cells, which could produce self-sustaining activity that supports a short-term memory trace. The distinct initial sensory cues, together with the behavior elicited by these cues, may push the network state of CA3 into one of two separate attractors representing right- and left-cued trials. As a result, CA3 cells will show differential responses later during the navigation even though the animal receives the same sensory inputs in the delay zone on the two trial types. Other synaptic pathways may also contribute to the context-dependent coding. For example, the nucleus reunions (NR) has been reported to deliver context-dependent information from the prefrontal cortex to CA1 (Ito *et al.*, 2015). In addition, context-dependent firing has also been described in the entorhinal cortex (EC) (Lipton, White and Eichenbaum, 2007), another major source of input for CA1. However, while it has been shown that CA3 inputs to CA1 can be potentiated by BTSP, whether NR and EC synapses can be potentiated similarly requires further investigation.

BTSP allows the conjunctive code in upstream circuits to be flexibly and accurately read out by CA1 cells. BTSP is induced by calcium plateau potentials, which require strong synaptic inputs on distal apical dendrites (Takahashi and Magee, 2009; Jarsky *et al.*, 2005). Therefore, layer 3 of the entorhinal cortex (EC3), which provides the most abundant excitatory distal inputs to CA1 pyramidal cells, serves as an instructive signal (Bittner *et al.*, 2015). What causes the activation of EC3 cells and thus the modification of CA1 representations remains unclear. Once plateau potentials are triggered, the resulting synaptic plasticity quickly transforms separate representations in the upstream areas into single-cell selectivity in CA1. With the somatic current injection protocol used in this study, plateau potentials in only a few trials can produce a stable large-amplitude place field, consistent with previous reports (Bittner *et al.*, 2015; Bittner *et al.*, 2017; Grienberger *et al.*, 2017; Zhao *et al.*, 2020). Importantly, under natural conditions, a single plateau potential is sufficient to produce a place field with the amplitude comparable to spontaneous place fields recorded in CA1 (Bittner *et al.*, 2015), likely due to the relatively large and prolonged local depolarization from the synaptically driven plateaus. Such single-trial induction ensures the selectivity of plasticity-induced response fields in CA1. Since the animal must be doing only one behavior at the moment when the induction happens, the consequence of such plasticity will be specific to that single behavioral context. In contrast, selectivity, even when high in the presynaptic area, will be averaged out in postsynaptic cells if the transformation is implemented through plasticity that requires a large number of induction trials, like spike timing-dependent plasticity (Bi and Poo, 1998; Markram *et al.*, 1997), because the animal can be engaged in different behavioral contexts during these trials.

The plasticity-induced response fields in the delay zone strongly depended on initial cues, but only after the animals were trained to associate these cues with reward locations. In animals trained with the random reward protocol, the plasticity induced nearly identical fields in trials with both cue configurations. This observation parallels previous reports that place cells differentiated trajectories within an environment only when rats were engaged in context-discriminating tasks (e.g., Gill, Mizumori and Smith, 2011). These results are consistent with our interpretation that separate attractors representing different cue configurations are formed in CA1’s upstream areas during training, which elicit distinct behavioral outcomes.

Our study also provides insights into the relative contributions of the past sensory cue and choice-related factors in plasticity-induced response fields by analyzing neural activity in error trials, where these two factors dissociated. Diverse responses were observed in CA1 cells, suggesting that two types of neural activity occur during errors: either the hippocampal neurons represent the trial-type information incorrectly, or the hippocampal trial-type encoding is apparently correct, but the animal’s behavior does not reflect a correct choice suggested by the hippocampal code, likely resulting from erroneous neural processes downstream of the hippocampus. Due to the low number of error trials in well-trained animals and the limited sample size from intracellular recordings, it is hard to quantitatively assess the prevalence of these two error trial types. High throughput recording or imaging methods could be used to address this issue in the future.

Our pharmacological silencing experiments reveal that the dorsal hippocampus is indispensable for animals to perform the cued Y-maze task in virtual reality. Since the silencing was conducted after the training, these results indicate that the hippocampus is required for the animal to retrieve learned rules to guide navigation, and/or support the short-term memory of initial cues during navigation. Previous studies have shown that, although context-dependent codes were observed in many behavioral paradigms, whether the hippocampus is required for these behaviors depended on nuances of the task design (Ainge *et al.*, 2007b; Wang *et al.*, 2015). The hippocampal dependence of our task may result from the relatively high cognitive demand for an animal to choose running directions based on visual cues with a randomized trial structure, compared to simply alternating directions from trial to trial, which rodents have intrinsic tendency to do (Lalonde, 2002). However, our results do not determine whether the context-dependent firing observed in the delay zone is specifically required for the behavior. First, it is unclear whether the hippocampus is only required during the initial cue zone to retrieve the memory of the task rules, or if it is required to correctly navigate the entire trajectory throughout the delay zone. While long-term memory of the rules is required for the cued Y-maze task, we do not know whether a short-term, working memory is also involved, as most mice had already exhibited differential behaviors in the cue zone. Future experiments using rapidly reversible optogenetics may be able to address this issue by silencing the hippocampus during particular task phases. Second, it is unclear whether the context-dependent firing, rather than the spatial tuning alone, is required to perform the task. Recently developed closed-loop optical approach (such as Robinson *et al.*, 2020), which enables manipulating functionally specific neuron ensembles, could be used to address the behavioral importance of splitter cells.

The hippocampus has long been considered to play a general role in memory-guided behaviors (Squire, 2009). Recent research is suggestive of a unified theory of hippocampal computations for both spatial and non-spatial memories (Aronov, Nevers and Tank, 2017; MacDonald *et al.*, 2013; Eichenbaum and Cohen, 2014). We expect that our findings are applicable not only to cued binary choice navigation but also to many other tasks where the hippocampus has been shown to exhibit a conjunctive code (MacDonald *et al.*, 2013; Sakon *et al.*, 2014). As we showed in the navigation task, slow learning could also establish conjunctive codes upstream of CA1 in non-spatial tasks, and the rapid plasticity induced by plateau potentials can flexibly and accurately route this code to other brain areas through CA1 to drive adaptive behaviors. Future work using chronic imaging will be essential to reveal the organizational principles underlying the emergence of context-dependent hippocampal maps during learning.

## Acknowledgements

We thank Mark Bolstad for technical help. We thank Dr. Albert Lee, Dr. Brett Mensh and all Spruston lab members for comments on the manuscript. This work is supported by Howard Hughes Medical Institute.

## Author Contributions

XZ, CLH and NS designed the project. XZ and CLH performed the experiments and analyzed the data. XZ, CLH and NS wrote the manuscript.

## Methods

### Animal Surgeries

Adult male mice (C57Bl/6, 6-12 postnatal weeks, Charles River Laboratories) were anesthetized with isoflurane (3-4% for induction and 1.5-2% for maintenance) and mounted on stereotaxic (Kopf Instruments) with ear bars. The scalp was removed, and the skull was cleaned and roughed by scalpels. Coordinates for craniotomies were measured by a micro-manipulator (MP285, Sutter Instrument) and marked with a fine marker pen. Two coordinates were marked for each hemisphere. One was to access CA1 (2.0mm posterior and 1.7mm lateral from Bregma) and the other (3.4mm posterior and 1.7mm lateral from Bregma) for the local field potential recording (LFP, see details in Electrophysiology). The skull was first covered with super glue. A customized titanium headbar was then implanted on the skull with Ortho-Jet dental cement (Lang Dental Manufacturing). The window of the headbar was covered by silicone sealant (Kwik-Cast, World Precision Instrument).

After the animals met the behavioral criteria after training, they were anesthetized and mounted on the stereotaxic again. Two small craniotomies (on one hemisphere), with diameters of 250-350μm, were made with a dental drill and covered with Kwik-Cast sealant. In some animals, the electrophysiology recordings started on the same day of the craniotomy surgery, with at least 2h recovery in the home cage. In other animals, the recordings started on the next day. The recordings continued for 2-4 consecutive days. Kwik-Cast was used to cover the craniotomy after each recording session. The same procedure was then repeated on the other hemisphere.

All experiments were performed according to protocols approved by the Janelia Research Campus Institutional Animal Care and Use Committee.

### Virtual Reality and Behavioral Training

The animals were head fixed on a virtual reality system as described before. The visual scene was rendered by Jovian, a VR engine developed at Janelia Research Campus, HHMI (Cohen, Bolstad and Lee, 2017; Zhao *et al.*, 2020), and displayed on 3 monitors (463UN, NEC) in front the animals that covers 216° in azimuth and ~90° in elevation. The optical flow was coupled with the animal’s locomotion on a spherical treadmill. Animals turned the spherical treadmill in three axes, pitch, roll and yaw. Throughout the study, animals were trained to move forward with pitch, while turn in a specific direction with roll. To prevent the animal’s running directions from causing differences in the visual input, the animal was lock at the midline of the track and its head direction remained straight before the joint point. The animal’s movement speed is determined by its velocity component projected to the forward direction. The animal’s ability to turn with the roll was released from the joint point. Animals ran uni-directionally in a Y-maze. At the end of each trial, the visual scene was frozen for a few seconds before the animal being teleported to the start location. The inter-trial-interval was 3 s for correct trials and 6s for incorrect trials. Animals collected a drop of sugar water reward (10% sucrose) when ran to the end of the correct arm. Licking was monitored by an infrared beam breaker (FX-300, Panasonic). Reward delivery and teleportation was programmed through a microcontroller (chipKIT Max32). A customized MATLAB code was used to record all behavioral variables at the frame rate (60Hz).

The animals entered water restriction schedule after ~1week recovery following the headbar surgery. Behavioral training started after ~1 week of water restriction. The animals received 1.5 ml water per day initially, and the daily water amount was adjusted (within the range of 1.0-2.0 ml) during the training based on the animal’s performance and body weight. The water amount was reduced if the animal showed low motivation (reluctant to run), while increased if the animal’s health deteriorated.

VR training followed a progressive curriculum (Supplementary Fig. 1). Various goals were set for each training stage. In maze A, the major goal was to train the animal get used to head fixation and run smoothly on the spherical treadmill. In maze B, the major goal was to let the animal learn to turn into both directions. All animals had an intrinsic gait bias with only running in a certain direction. The animal would stick at the blocking wall in maze B when its innate preference mismatched the cue direction until it switched to the other direction. After a few days training in maze B, the animal turned in both directions before hitting the blocking wall, which indicated that the animal was ready to be moved on to maze C. The animal was trained in maze C until it performed at >75% success rate. The animal was then trained in maze D, E, and F, sequentially, to learn to perform with the delay period and a longer track. The animal was moved back to the previous stage if its performance dropped and did not show improvement in a few days when new maze was introduced. All animal reached success rates higher than 80% in the final maze (maze F) before physiological or behavioral experiments began. The whole training process typically took 3-6 weeks.

One challenge for mice to perform the cued Y-maze task is that they exhibited biased turning directions when running on the spherical treadmill. A few approaches were used to help animals overcome this strong bias. First, walls were used to block the wrong side, as described before. Second, during the initial stage of training without blocking walls, an error-trial-repeating mechanism was introduced. For example, if the animal turned into the left arm in a right-cued trial, the probability of the next trial being the right-cued trial was increased to 67-80%. It should be noted that such error-trial-repeating mechanism was turned off finally in all physiological and behavioral experiments. Third, in some animals a large block of trials (tens of trials) with their non-preferred directions were used in the early phase of training.

The VR gain was set high (2-3) initially to encourage running. The gain was gradually reduced through training. It was reduced eventually to make sure the peak running rate was approximately 25-30 cm/s. The VR gain was not changed in the final phase of training (maze) and during the experiments.

In the random reward group, sugar water rewards were delivered at 75-90% of chance, regardless of the initial cue.

To assist balanced sampling of left- and right-cued trials during limited time of a recording session, at most 4 consecutive trials with the same cue direction was allowed. In another word, if 4 trials were already cued to the same direction, the next trial would be cued to the other direction. Cue directions were randomized in all other situations.

### Pharmacology and Histology

BODIPY TMR-X muscimol conjugate (Invitrogen) was used to silence hippocampus. TMR-X muscimol conjugate was first dissolved in DMSO at 16mM and then diluted by 5-fold in 0.9% saline. Some deposit was observed during the dilution. The solution was vibrated at 45°C, 1000rpm to facilitate dissolution. The final concentration is 3.2 mM muscimol in 20% DMSO.

Three animals used in the silencing experiments (Fig. 1) were animals previously used for physiological recordings. The other three animals were used only for the silencing experiments. The performance in the Y-maze task was first tested to ensure that the animal performed at >80% success rate. On the next day, animals were anesthetized and mounted on the stereotaxic. For silencing-experiment-only animals, craniotomies on top of hippocampus on both hemispheres were drilled. The procedure took approximately 1h. For animals previously used for recordings, they were also anesthetized for 1h with the surgical lamp on throughout the whole time. The previously prepared craniotomies were cleaned by fine forceps since sometimes tissue regrowth occurred. The animal was then taken off anesthesia and then recovered in its home cage for 1h, before transferred to the VR rig for the behavioral test. This test is to confirm that 1h recovery is enough for the animal to recover from potential adversary effects of anesthesia and strong light on its ability to behave in VR. On the next day, the animal was anesthetized again. 3.2mM TMR-X muscimol was injected bilaterally into dorsal hippocampus through Nanoject (Drummond Scientific). Per one hemisphere, injections were made at 2 depths (1.25 mm and 1.65 from pia), 27.6 nl per depth. The injection procedure typically took ~1h. The craniotomies were covered by Kwik-Cast, and the animal was allowed to recover for 1h before the behavioral test.

Three animals in the silencing experiment were euthanized by overdosing isoflurane immediately after the behavioral test. The animals then underwent trans-cardiac perfusion with 4% parahydroxyamide (PFA). The brain was dissected out and further fixed in 4% PFA for overnight. Another three animals were saved for one more day for another behavioral test to examine the reversibility of muscimol’s effect. These three animals were euthanized and perfused after the second test.

Coronal sections, 100μm thick, were cut from fixed brain samples using a vibrotome (Leica). Brain sections were mounted on coated glass slides and covered with mounting media containing DAPI (VectaShield, Vector Laboratories). Images of brain slices were taken using a confocal microscope (Zeiss 880 Airyscan).

### Electrophysiology Recording

Whole-cell recordings were performed in CA1 pyramidal layer using standard blind patching technique, as described before (Zhao *et al.*, 2020). Glass pipettes were made with P-97 puller (Sutter Instruments). The pipette impedance was 10-13MΩ. The glass pipette was inserted vertically using a micro-manipulator (4-axis *in-vivo* unit, Luigs and Neumann) into the hippocampus. The pipette was filled with internal solution containing (in mM): 134 K-gluconate, 6.6 KCl, 10 HEPES, 3.4 NaCl, 0.3 Na_2_-GTP, 4 Mg-ATP and 14 Tris-phosphocreatine (all from Sigma). A second glass pipette (1-3 MΩ), filled with 0.9% saline, was inserted from the posterior craniotomy at 45° into the hippocampus with a hydraulic manipulator (Narishige) to monitor LFP. This pipette also served to stabilize the patch-clamp recording. A chloridized silver wire was placed in the headbar window as the ground. The headbar window was covered with 0.9% saline. The patching pipette was initially advanced into CA1 (before reaching the pyramidal layer) with a large positive pressure (6-8psi). The pressure was then reduced to ~0.35 psi and the cell searching began. Standard procedure was followed for sealing and breaking in the cell. Current clamp recordings were conducted in all experiments. The identity of pyramidal cells was confirmed by their firing pattern. As shown in previous *in-vivo* whole-cell recordings, the cell appeared very quiet within a few minutes after the break-in, potentially due to the puffed out high potassium internal solution during patching. Recordings started only after normal spontaneous activities came back.

Electrical signals were amplified with BVC-700A amplifier (Dagan) and recorded through NI USB-6343 acquisition board (National Instruments). Data were digitized at 20 kHz. Capacitance compensation and bridge balance were done manually. Series resistance (R_s_) was determined from the bridge balance. Only cells with Rs < 100 MOhm were included in our analysis. In some cells, small hyperpolarizing holding currents (<100 pA) were applied to maintain the cell’s membrane potential. The holding current was not changed during the recording. To induce BTSP, 300 ms square pulse current was injected once the animal ran to a certain location. The amplitude of the current injection was adjusted at 700-900 pA, determined by the experimenter to ensure the occurrence of plateau potentials. To synchronize behavior and electrophysiology data streams, 1 Hz TTL pulses were generated by the acquisition board and recorded by both the board itself and the micro-controller used to control the behavior system. WaveSurfer, a MATLAB based software developed at Janelia Research Campus, was used to record electrophysiology data. Junction potential was not corrected in all reported numbers.

### Data Analysis

Electrophysiological recordings were first aligned with behavioral data using sync-pulses. Left- and right-cued trials were sorted, and the position at each time point was calculated as the distance from the start location. The distance data were then interpolated to match the high sampling rate of physiological recordings (20k).

Hippocampus is known to have distinct states during stationary and locomotion periods. In this study, only data within running periods (speed>2cm/s) were used in characterizing cellular activities.

To analyze the firing rate, dV_m_/dt was calculated, and the initiation of each action potential was identified as dV_m_/dt > 3 mV/s. In a few recordings with noisy conditions, dV_m_/dt was first filtered with a low-pass filter (0-1000Hz) and dV_m_/dt(filtered) > 0.4 mV/s was used as the criterion. To calculate the spatial tuning of firing rates, the whole track was binned by 2.5 cm. The firing rate in each bin was calculated as the number of action potential divided by the occupancy time. The peak firing rate was defined as the maximal firing rate in the delay zone, for each trial type. Peak rate difference between the two trial types was calculated as the difference between the two rates divided by the larger one.

For all analyses of subthreshold Vm dynamics, action potentials were first removed by a 10 ms median filter.

Ramp V_m_ was calculated by applying a 0-2 Hz low-pass filter. The spatial tuning curve of ramp V_m_ was calculated by averaging ramp Vm within 0.1 cm spatial bins. Ramp depolarization was then calculated by subtracting the averaged ramp Vm within the first 1cm from the ramp curve. Ramp depolarization was averaged across trials for each trial type. Finally, the ramp depolarization curve was smoothened by a Gaussian filter with the S.D. of 12.5 cm. Position was found where the maximal ramp depolarization occurred in plasticity trial. Averaged ramp depolarization was calculated within a +/-5 cm window around the peak location as the peak ramp depolarization. Peak ramp depolarization for the opposite trial was calculated from the same window.

Baseline Vm was defined as the average of 10 minimal ramp Vm data points (corresponding to 1cm), without any subtraction.

Theta Vm was calculated by applying a 4-10 Hz band-pass window. The amplitude of theta oscillation was calculated by the Hilbert transform of the theta Vm. The spatial map of theta amplitude was quantified and smoothened in the same way as described above for the ramp Vm.

To analyze the spatial profile of running speed, pitch and roll data were interpolated at 20k and averaged across trials, for each trial type. Yaw was not analyzed since it was not couple with VR rendering. The spatial map of running speed is smoothened by a Gaussian filter with 12.5 cm S.D. For the lateral running speed, positive values indicate left turning, while negative ones indicate right turning. The total locomotion speed was calculated as:

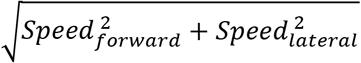

All statistical tests were conducted with two-tailed paired student t-test. P<0.05 was used as the criterion for significance. In all curve plots, curves and shades represented mean and S.E.M., respectively. Normal distribution of the data was assumed, as in previous studies with similar methodology, but not rigorously tested. In all box-and-whisker plots, center lines depict the median; boxes depict 1 inter quartile range (IQR); and whiskers go to the largest/smallest data point within the ±1.5 X IQR range. Note: the whisker range of 1.5 X IQR was just chosen for depiction, but all data points were included in statistical comparisons. No points were removed as outliers. All analyses were done with customized MATLAB codes (version 2018, MathWorks).

