## Supplemental Figures for "Rapid synaptic plasticity contributes to a learned conjunctive code of position and choice-related information in the hippocampus"

| Cell # | Cued Direction during the Induction | Correct (C) or Error (E) Trial during the Induction | % Peak Rate Difference | Classified as Splitter Cells |
|---|---|---|---|---|
| 1 | Left | C E E C C | 23.7 | No |
| 2 | Right | C E C C | 14.8 | No |
| 3 | Right | C | 96.2 | Yes |
| 4 | Left | C C C C | 100 | Yes |
| 5 | Left | E C C | 100 | Yes |
| 6 | Right | E C C | 95.9 | Yes |
| 7 | Left | C C C | 100 | Yes |
| 8 | Right | C C | 93.9 | Yes |
| 9 | Right | C C C C C | 100 | Yes |
| 10 | Left | E E C C | 81.7 | Yes |
| 11 | Right | C C C C C | 100 | Yes |
| 12 | Left | C C C C C C | 93.0 | Yes |
| 13 | Left | C C E C | 90.5 | Yes |
| 14 | Left | C C C | 71.0 | Yes |

**Supplementary Table 1. Induction of place fields by experimentally triggered plateau potentials.** All cells included in Fig. 2 are listed. Correct (C) and error (E) trials are labeled in green and red, respectively.

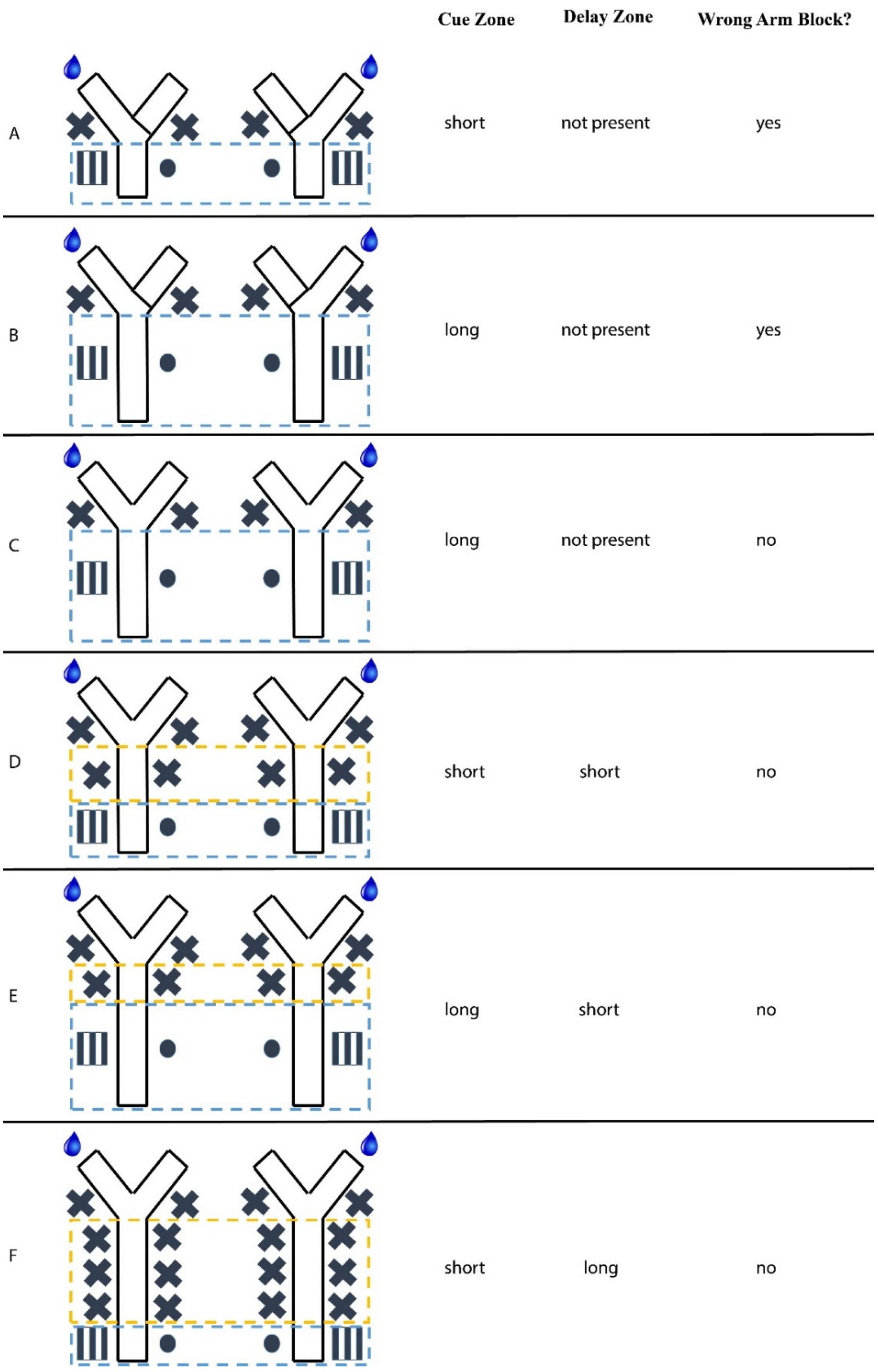


**Supplementary Figure 1. Progressive training curriculum.**

Each animal was trained through A to F, with F being the final task where all behavioral and physiological experiments with fixed cue-reward association were conducted. Overall, as the task difficulty increased, the mouse's performance enhanced, but in some cases the performance was not stable or did not continue to improve, an easier maze could be used again. Blue and yellow boxes depict cue and delay zones, respectively. Water droplet represents the rewarded location for a given trial.

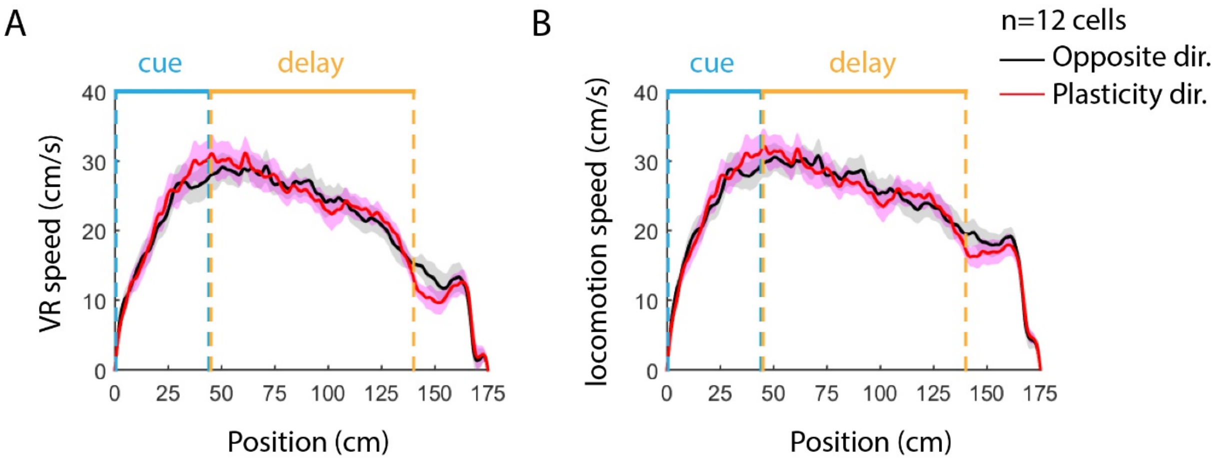


**Supplementary Figure 2. The speed of VR optic flow (A) or locomotion speed of the animals (B) did not differ between trials of the plasticity vs. opposite directions.**

Curves and shades represent mean and S.E.M (red: plasticity direction, black: opposite direction, n=12 cells from 6 mice). Blue and yellow bars mark the cue and delay zones, respectively. The VR speed was calculated from the animal's forward running speed. The locomotion speed was defined as the vector length of the animal's true running velocity, independent of its direction. The VR speed is the projection of the locomotion speed along the forward direction of the VR environment.

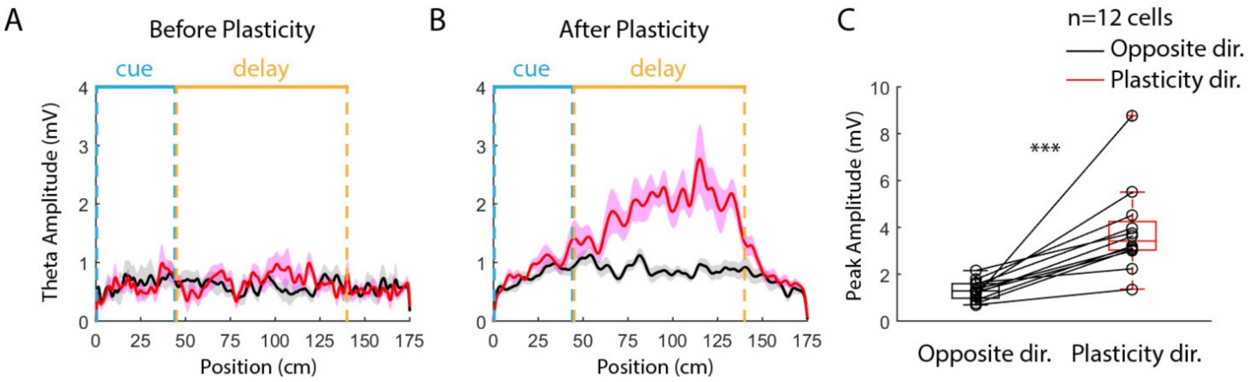


**Supplementary Figure 3. Plasticity-induced context-dependent place fields exhibited enhanced theta oscillation.**

**A and B.** Averaged theta amplitude before (A) and after (B) the plasticity for induced splitter cells (red: plasticity direction, black: opposite direction, n=12 cells from 6 mice). Curves and shades represent mean and S.E.M. Blue and yellow bars mark the cue and delay zones, respectively.

**C.** Statistical comparison of the peak theta amplitude in the delay zone between trial types (p=6.795e-4; two-tailed paired student t-test; n=12 cells from 6 mice). In the whisker plot, center lines depict the median; boxes depict 1 inter-quartile range (IQR); whiskers indicate the largest/smallest data value within the ±1.5 X IQR range. Note: all data points were included in statistical comparisons, with no points removed as outliers.

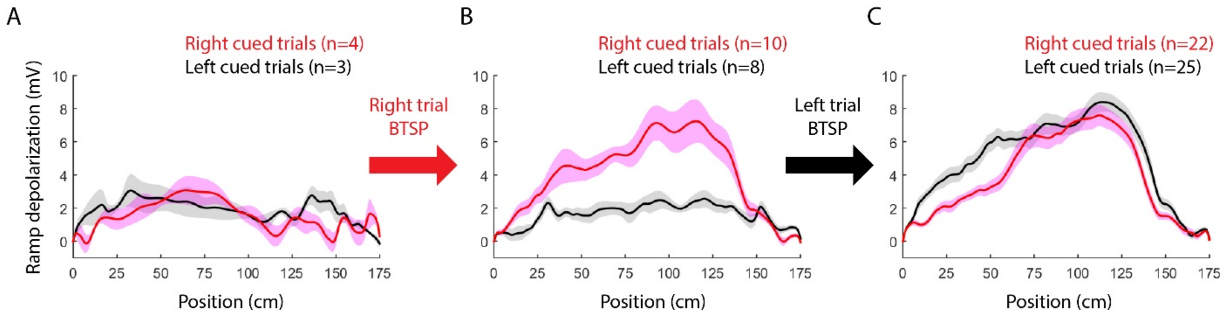


**Supplementary Figure 4. Consecutive inductions of two trial-type specific place fields in one cell.**

Ramp depolarizations were plotted against the animal's position. Curves and shades represent mean and S.E.M., respectively. n represents the number of trials.

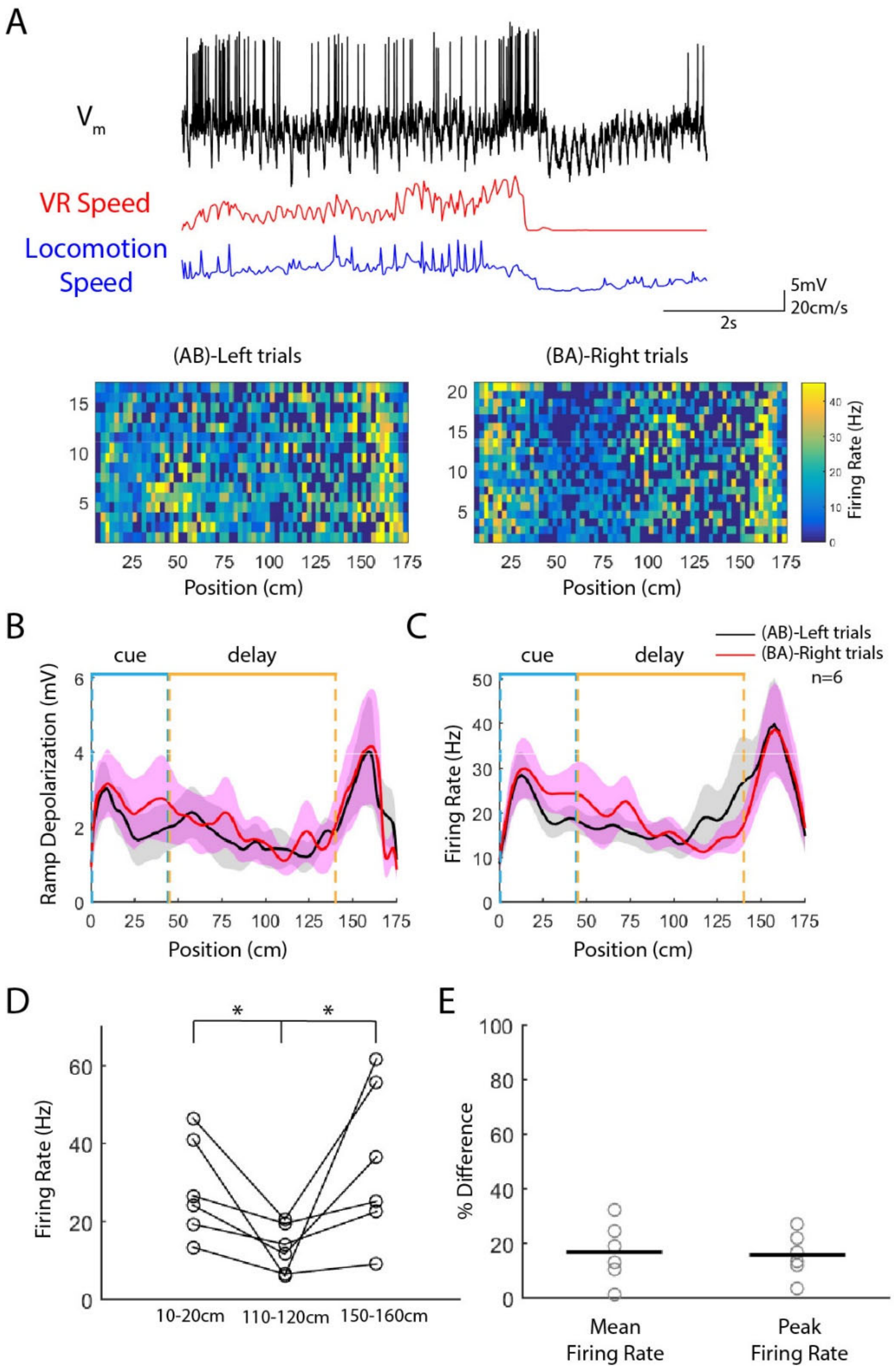


**Supplementary Figure 5. Putative fast-spiking interneurons were only moderately modulated by trial type.**

**A.** Example recording of a putative fast-spiking interneuron. The cell had a high firing rate and was highly modulated by the animal's running (locomotion) speed. There was a period of the time near the end of the trial when the VR speed, but not the locomotion speed, was zero because the mouse was at the end of the virtual track before being teleported. *Bottom*: trial-by-trial spatial heat maps of firing rate sorted by trial type (spatial bin: 2.5cm).

**B.** Ramp depolarization of putative fast-spiking interneurons in left- (black) vs. right-cued (red) trials. Curves and shades represent mean and S.E.M, respectively. n=6 cells from 3 mice.

**C.** Firing rate of putative fast-spiking interneurons in left- (black) vs. right-cued (red) trials. Curves and shades represent mean and S.E.M, respectively. n=6 cells from 3 mice.

**D.** Averaged firing rate at select locations within the cue (10-20cm), delay (110-120cm), and reward (150-160cm) zone. Recordings from both trial types were pooled together for the measurement. P=0.0273 for cue vs. delay, 0.0477 for delay vs. reward (paired Student's t-test). n=6 cells from 3 mice.

**E.** Normalized difference in the mean and peak firing rates in the delay zone between two trial types. Black ticks represent mean value (16.8% for the mean rate and 15.8% for the peak rate). n=6 cells from 3 mice.

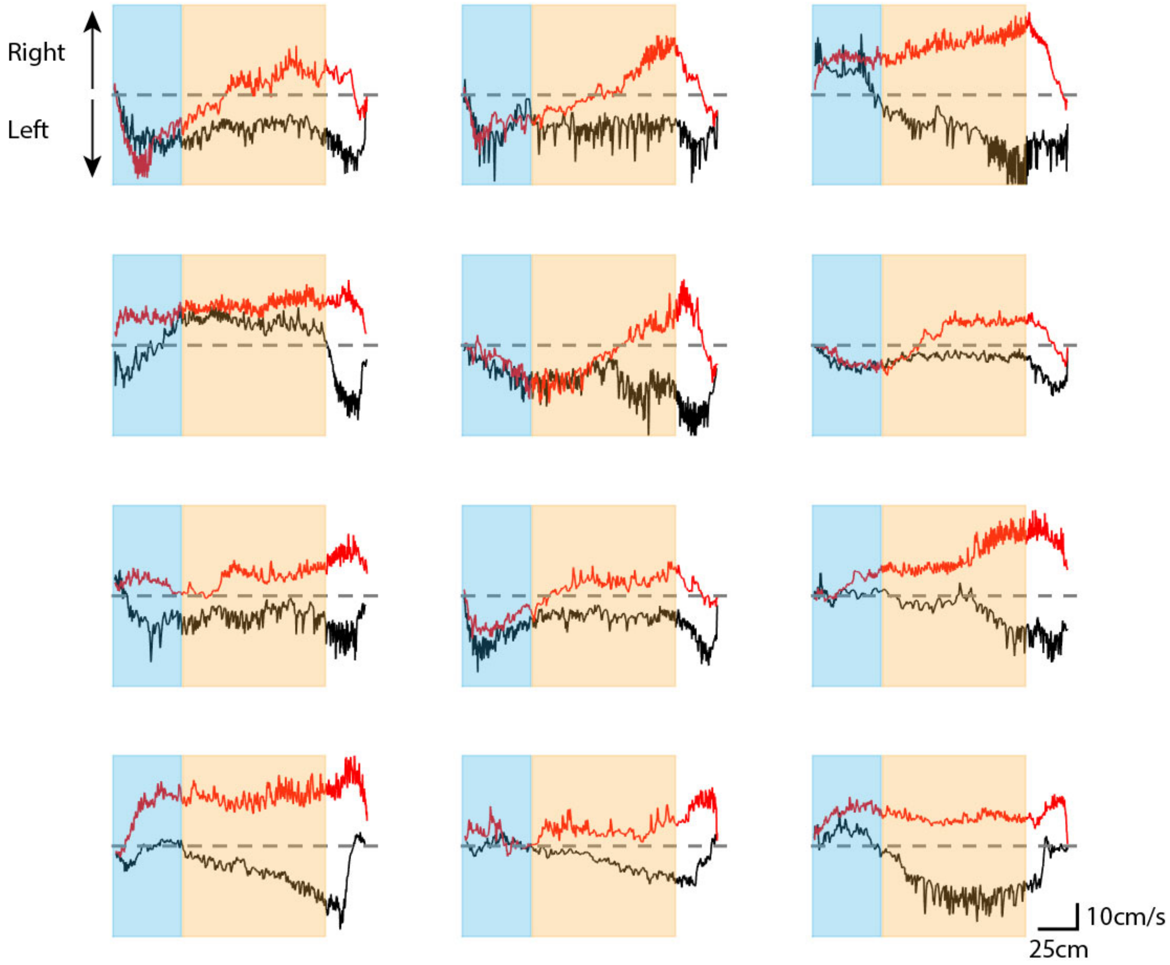


**Supplementary Figure 6. Lateral running speed for all 12 induced splitter cells.**

Positive values indicate right turning, while negative values indicate left turning (zeros, indicating the midline, were depicted in grey dashed lines). Red: right-turning trials; black: left-turning trials. Blue and yellow shades represent cue and delay zones, respectively. Curves were averaged across trials for each cell.

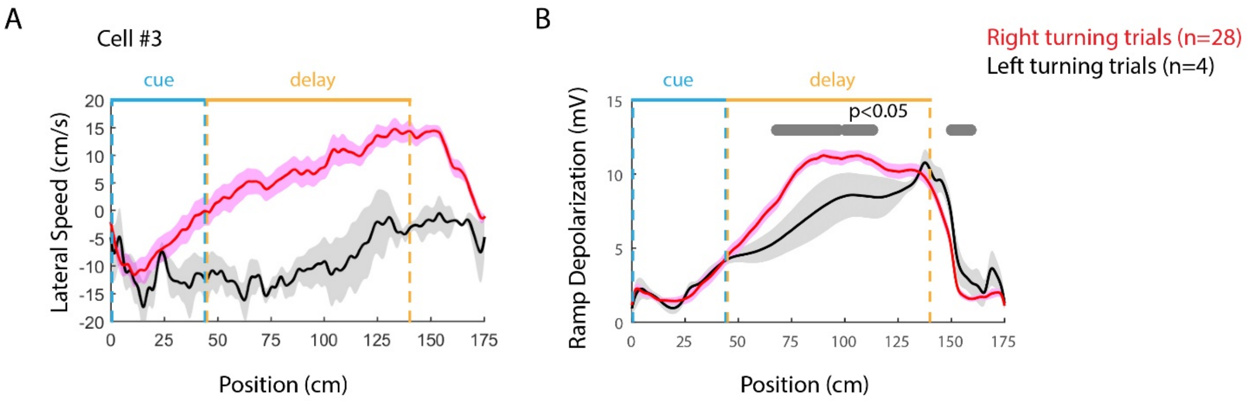


**Supplementary Figure 7. Running direction modulated the place-field activity only moderately in one animal trained with the random reward protocol.**

Lateral running speed (A) and ramp depolarization (B) were plotted in right- (red) and left-turning (black) trials. Curves and shades represent mean and S.E.M, respectively. Grey bars in B mark the positions where the red and black curves are significantly different ($p<0.05$, two-tailed paired Student's t-test). In this cell, the mouse turned to the right side during all of the induction trials when plateau potentials were triggered. n represents the number of trials.

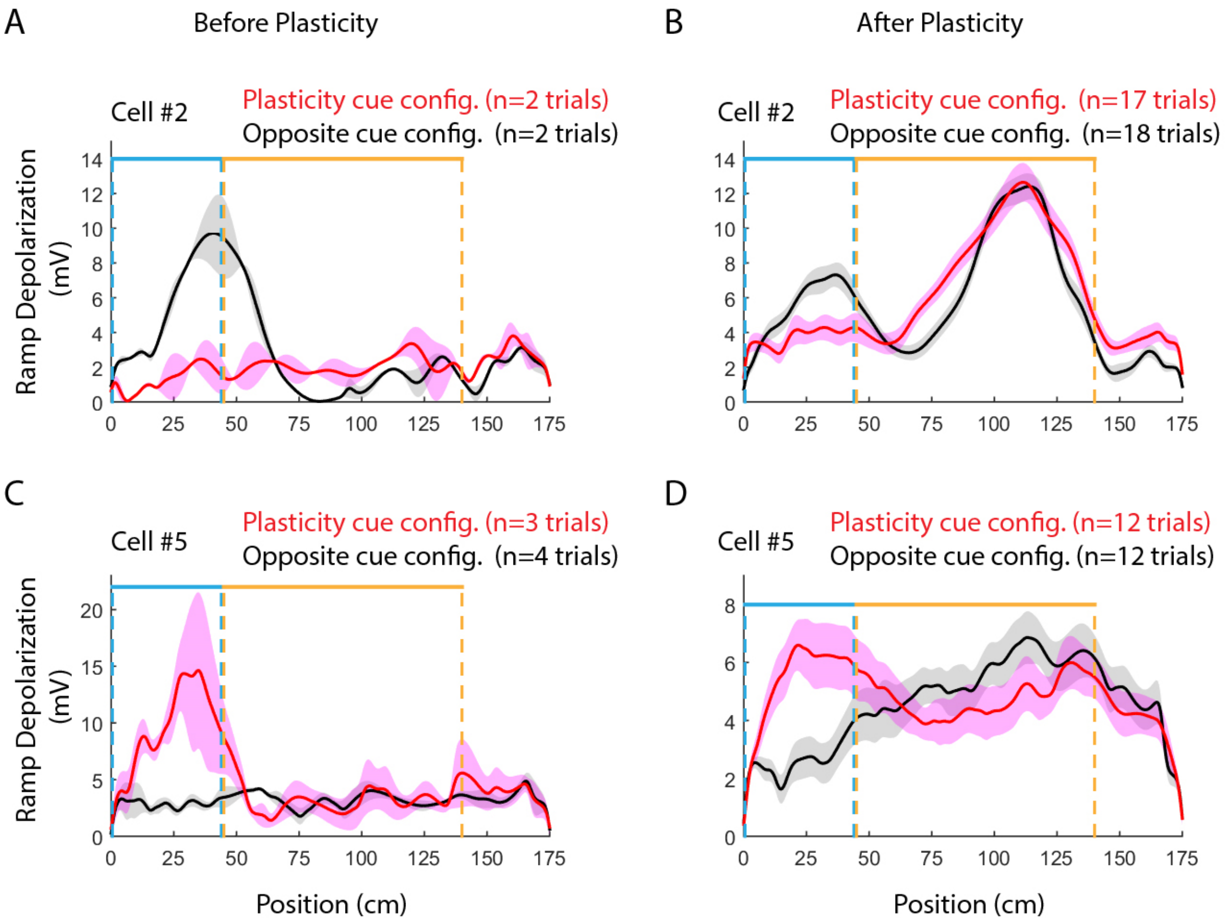


**Supplementary Figure 8. Distinct immediate visual cues in the cue zone drove different place fields in two CA1 cells from animals trained with the random reward protocol.**

Ramp depolarization was plotted against position for right- (red) and left-cued (black) trials from two cells which exhibited native place fields prior to plasticity induction. These cells were from the animals trained with the random reward protocol. Place fields starting in the cue zone were trial-type-specific (i.e., depended on the initial cues) before (A and C) and after (B and D) the plasticity induction. Curves and shades represent mean and S.E.M, respectively.
